# GIGGLE: a search engine for large-scale integrated genome analysis

**DOI:** 10.1101/157735

**Authors:** Ryan M. Layer, Brent S. Pedersen, Tonya DiSera, Gabor T. Marth, Jason Gertz, Aaron R. Quinlan

## Abstract

GIGGLE is a genomics search engine that identifies and ranks the significance of shared genomic loci between query features and thousands of genome interval files. GIGGLE scales to billions of intervals, is faster (+1,000X) than existing methods, and its speed extends the accessibility and utility of resources such as ENCODE, Roadmap Epigenomics, and GTEX by facilitating data integration and hypothesis generation. GIGGLE is available at https://github.com/ryanlayer/giggle.

## INTRODUCTION

The results from genome-wide assays (e.g., ChIP-seq, RNA-seq, variant calling) are often interpreted by comparing experimentally identified genomic loci to other known genomic features (e.g., open chromatin, enhancers, transcribed regions). While large-scale functional genomics projects have greatly empowered this type of analysis by characterizing the genomic regions associated with a wide range of genomic processes, interpretation is complicated by the thousands of results that span hundreds of different tissue types, assays, and biologic conditions. Effectively integrating these large, complex, and heterogeneous resources requires the ability to rapidly search the full dataset and identify the most statistically relevant features. While existing software such as BEDTOOLS^1^ and TABIX^2^ identify regions that are common to genome interval files, these methods were designed to investigate a limited number of files. More recent methods^3, 4^ describe improved statistical measures, yet do not scale to the vast amount of data that is now available.

We introduce GIGGLE as a fast and highly scalable genomic interval searching strategy that, much like web search engines did for the Internet, provides users with the ability to conduct large-scale comparisons of their results with thousands of reference datasets and genome annotations in seconds. GIGGLE enables the identification of novel and unexpected relationships among local datasets as well as the vast amount of publicly available genomics data through command line and web interfaces, as well as APIs in the C, Go, and Python programming languages. GIGGLE is based on a temporal indexing scheme^5^ that uses a B+tree to create a single index of the genome intervals from thousands of annotations and genomic data files (Figure 1a). Each interval in an indexed file is represented by two keys in the tree that correspond to the interval’s bounds (start and end+1). Each key in a leaf node contains a list of intervals that either start at (indicted by a “+”) or have ended (indicated by a “-”) just prior to that chromosomal position. For example, in Figure 1a position 7 corresponds to a key in the second leaf node with the list [+T2,-B2]. This indicates that at chromosomal position 7, the second interval in the “Transcripts” file (T2) starts and the second interval in the “TF binding sites” file (B2) has ended. To find the intervals in the index that intersect a query interval (e.g. [1,5] in Figure 1a), the tree is searched for the query start and end (shown in bold red in Figure 1a), the keys within that range (shown in red in Figure 1a) are scanned, and intervals in the lists of those keys are identified as intersecting the query interval (see **Online Methods** for complete algorithmic details).

GIGGLE’s potential for high scalability is based on two factors. First, identifying the number of overlaps between a query and any given annotation file is determined entirely within the unified index, thus eliminating the inefficiencies of existing methods, which must instead open and inspect the underlying data files. Second, the B+tree structure minimizes disk reads; this is vital to performance since databases of this scale will grow beyond the capacity of main memory and must be stored on disk. To measure GIGGLE’s query performance, we created an index of the ChromHMM^6^ annotations curated by the Roadmap Epigenomics Project (Roadmap) from 127 tissues and cell lines. Each genome was segmented into 15 genomic states, yielding over 55 million intervals in the resulting GIGGLE index. When testing query performance with a range of 10 to 1,000,000 query intervals, a GIGGLE was 2,336X faster than TABIX and 25X faster than BEDTOOLS (Figure 1b) for the largest comparison. Similarly, using an index of 5,603 annotation files for the human genome (hg19, a total of 6.9 billion intervals) from the UCSC Genome browser, GIGGLE was up to 345X faster than TABIX and 8X faster than BEDTOOLS (Figure 1c).

**Figure 1.**
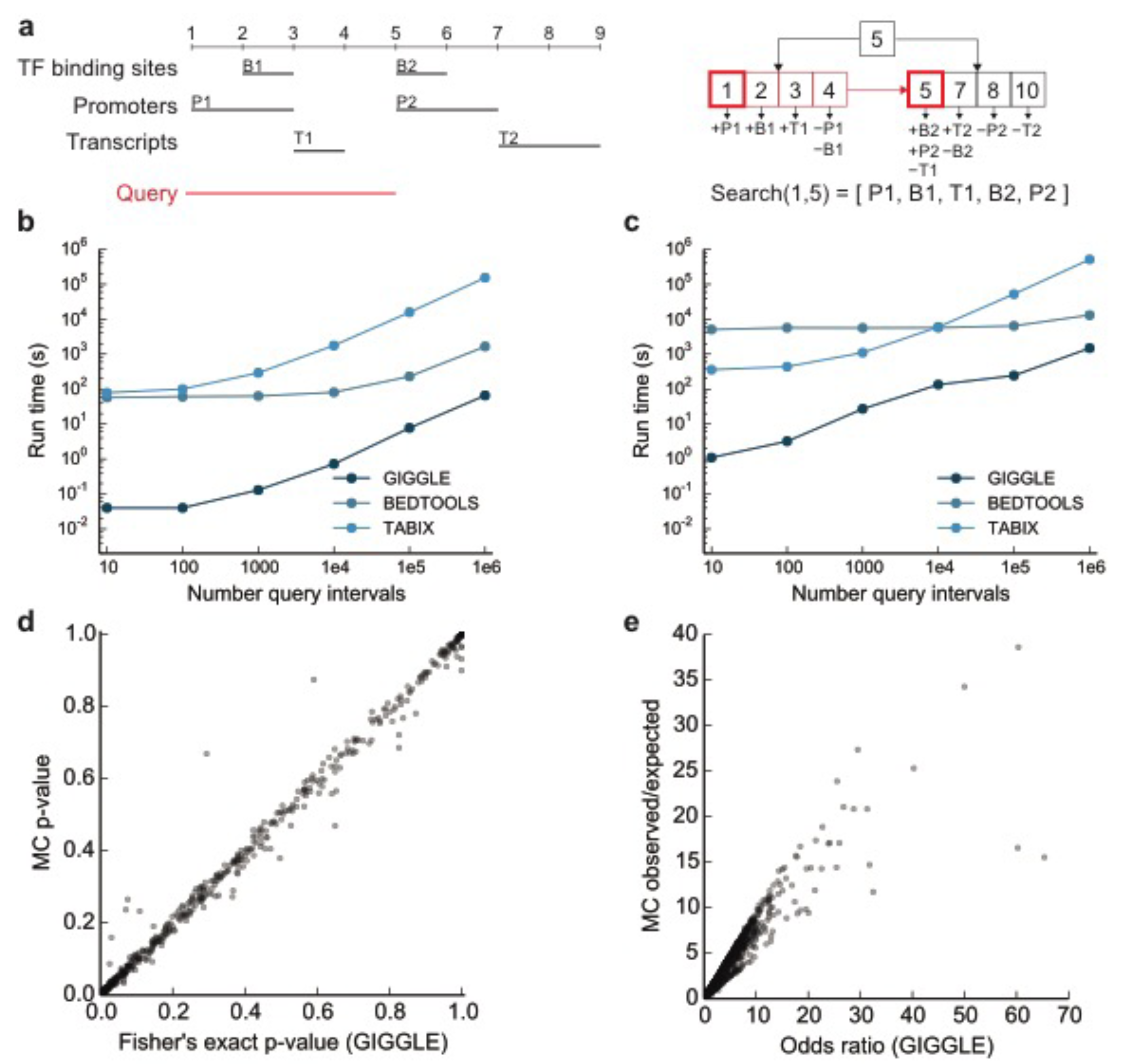
GIGGLE indexing, searching, performance, and score calibration. (**a**) A set of three genomic intervals files (transcription factor (TF) binding sites, promoters, and transcripts) (left, black) is indexed using a single (simplified) B+ tree (right). Intervals among the annotations overlapping a query interval (left, red) are found by searching the tree for the query start and end (right, bold red) and scanning the keys between these positions (right, red). (**b**) Run times for GIGGLE, BEDTOOLS, and TABIX considering random query sets with between 10 and 1 million random 100 base pair intervals against the CHROMHMM processing of Roadmap Epigenomics (1,905 files and over 55 million intervals). (**c**) Run times for the same method and queries against UCSC genome browser annotations (5,603 files and over 6.9 billion intervals). Here, the runtimes for GIGGLE and BEDTOOLS converge for query sizes exceeding hundreds of millions of intervals. While this is expected given the underlying computational complexities of the methods, this scenario far exceeds typical query set sizes. (**d,e**) A comparison between GIGGLE’s relationship estimates using a contingency table and Monte Carlo-based methods for (**d**) significance and (**e**) enrichment considering a search of MyoD ChIP-seq peaks against CHROMHMM predictions from Roadmap.

Speed is essential for searching data of this scale, but, as with internet searches, it is arguably more important to rank order results by their relevance to the set of query intervals. Ranking requires a metric that quantifies the degree of similarity between the query intervals and each interval file in the GIGGLE index. Monte Carlo (MC) simulations are commonly used in genomics analyses^7,8^ to compare the observed number of intersections to a null distribution of intersections obtained by randomly shuffling intervals thousands of times and testing the number of intersections in each trial. While MC simulations are an effective method for pairs of interval sets, they are computationally intractable for large-scale datasets since thousands of permutations are required for each interval file.

GIGGLE eliminates this complexity by estimating the significance and enrichment between the query intervals and each indexed interval file with a Fisher’s Exact Test and the odds ratio of a 2×2 contingency table containing the number of intervals that are in (1) both the query and indexed file, (2) solely the query file, (3) solely the indexed file, and (4) neither the query file nor the indexed file. The first three values are directly computed with a GIGGLE search, and the last value is estimated by the difference between the union of the two sets and the quotient of the mean interval size of both sets and the genome size. These estimates are well correlated with the Monte Carlo results (Figure 1d, Figure 1e) and have the favorable property of near instant computation.

GIGGLE ranks query results by a composite of the product of −log_10_(p-value) and log_2_(odds ratio). This “GIGGLE score” avoids some of the issues that arise when using only p-values to select top hits. In MC simulations, the proportion of values that are more extreme than the observation (i.e., the p-value) is highly dependent on the variance of the trials. When the variance MC distribution is low, observations that are only marginally larger than the expected value may be significant, yet not interesting biologically. For example, one result from the search of MyoD ChIP-seq peaks against Roadmap had a low enrichment (1.7X), but the variance of the MC simulations was also low (88.36) making the observation significant (p < 0.001). Similarly, when the MC distribution variance is high, large enrichments may not reach significance. These effects are mitigated by combining significance and enrichment into the GIGGLE score.

While the GIGGLE score can be used to rank results, it is also insightful to use all scores to visualize the full spectrum of relationships between a query set and all indexed interval sets. For example, a heatmap of GIGGLE scores from a search of MyoD ChIP-seq peaks against the GIGGLE index of Roadmap (Figure 2a) illustrates MyoD’s important and specific role in muscle tissues. Similarly, a search of GWAS SNPs associated with Crohn’s disease^11^ (Figure 2b) shows that variants cluster in immune cell enhancers. While these results illuminate the dynamics of individual features, the speed of GIGGLE (<0.3s for Crohn’s SNPs) allows researchers to conduct exploratory research on a massive scale. For example, GIGGLE can quickly (3.5s) query sets of GWAS SNPs for 39 different traits^11^ against Roadmap to not only confirm that the enrichment of SNPs in immune cell enhancers is present in other autoimmune diseases and absent in non-autoimmune traits^11^ (Figure 2c, left), but to also show that there is no cell-specific pattern in transcribed regions for either set of traits (Figure 2c, right).

We emphasize that GIGGLE is a completely general framework that enables researchers to efficiently explore any collection of intervals sets. For example, using a GIGGLE index of all ChIP-seq datasets available from Cistrome^12^ (5,992 files, 8,716,024 intervals), we quickly (<3 min) performed a full pair-wise comparison of the 270 factors (734,249 intervals) available for the MCF-7 breast cancer cell line. From this comparison, distinct subsets become clear, including coordinated genomic binding of CTCF, RAD21, and STAG1, indicative of regions involved in long-range interactions^13–15^, and estrogen receptor a (ER) co-occurrence with other transcription factors (Group 1 and Group 2 in Supplemental Figure 2, respectively). Focusing specifically on ER (Figure 2d) uncovers sequence-specific transcription factors known to play important roles in ER genomic binding (FOXA1^16^, GATA3^17^ and PR^18^) and co-factors that are involved in estrogen-induced gene regulation (EP300^19^ and NCAPG^20^). One unexpected finding from this large-scale analysis of MCF-7 ChIP-seq data is the strong co-occurrence of histone variant H2AFX^20, 21^ and ER co-factor GREB1^22^ (Group 3 in Supplemental Figure 2), suggesting a potential physical interaction between these factors.

GIGGLE also provides the infrastructure to integrate data sources. For example, we developed a web interface that allows users to further investigate interesting results from Roadmap (e.g., MyoD ChIP-seq and Myoblast enhancers) by a querying a GIGGLE index of the UCSC genome browser data (Supplemental Figure 3). Those results are visualized in the genome browser as a dynamic “smartview” where only the tracks with at least one overlap are visible. Other GIGGLE indices can also be used to verify results. For example, we recapitulated the top hits from the GIGGLE search of both MyoD ChIP-seq peaks and Crohn’s disease GWAS SNPs against Roadmap with similar searches against an index of the FANTOM5 data^23^ (1,825 files, 11,284,790 intervals) (Supplemental Tables 1–4). This is especially promising since FANTOM5 and Roadmap are based on fundamentally different assays, and therefore provide orthogonal corroboration of these biological relationships. These examples illustrate GIGGLE’s ability to confirm previously characterized associations, and demonstrate the discovery potential afforded by GIGGLE’s rapid, prioritized searches.

The exploratory power of a single interface from which many datasets can be searched is unmatched by any current method, and has the possibility to dramatically advance large-scale, integrative analyses. GIGGLE is capable of powering a single access point that will inform researchers and clinicians of all known experiments and curated annotations that are associated with a particular genomic region. In summary, GIGGLE provides a new engine with which to conduct large-scale, in silico “screens” of multi-dimensional genomics datasets in search of insights into genome biology in diverse experimental contexts.

**Figure 2.**
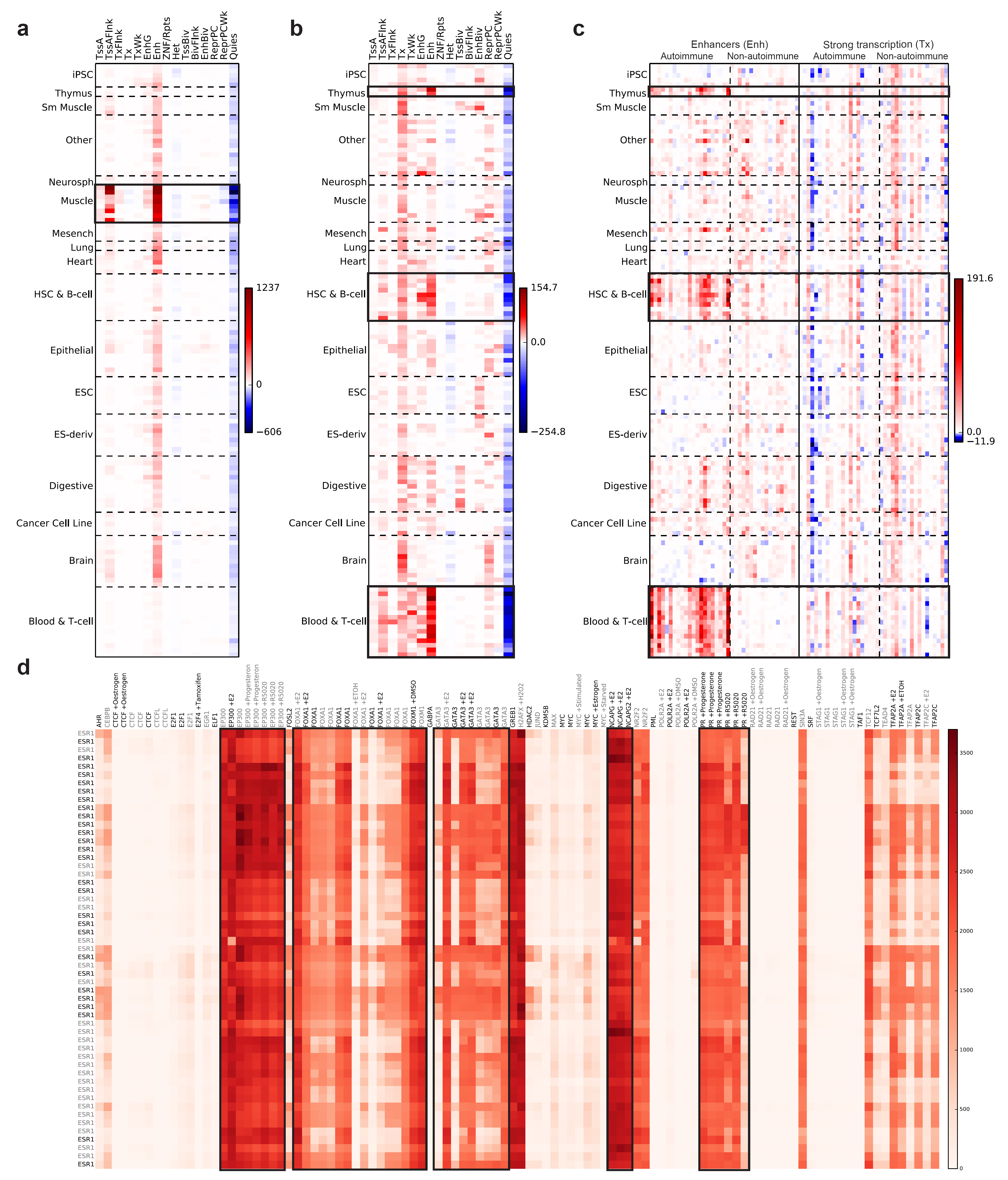
Visualization of GIGGLE scores from various searches. Darker red (higher GIGGLE scores) indicate more enrichment and darker blue (lower GIGGLE scores) indicate more depletion versus the expected value. The relationships between (a) MyoD ChIP-seq peaks and (b) genome-wide significant SNPs for Crohn’s disease and 15 genomic states across 127 different cell types and tissues predicted by ChromHMM for Roadmap. Black boxes within panels highlight (a) muscle and (b) immune tissues and cell types. (c) Results from the enhancer (major left column) and strong transcription (major right column) tracks from ChromHMM for Roadmap when considering GWAS SNPs for 21 autoimmune disorder (minor left) and 18 non-autoimmune traits (minor right). The black boxes highlight immune tissues and cell types. (d) The relationships between ESR1 ChiP-seq binding sites and the binding sites from 105 other ChiP-seq experiments (38 different unique factors) in MCF-7 cells. Darker red (higher GIGGLE scores) indicate more enrichment. Black boxes highlight the relationships between ESR1 and FOXA1, GATA3, PR, EP300 and NCAPG.

## FUNDING

This research was funded by NIH grants to RML (K99HG009532) and ARQ (R01HG006693).

## ONLINE METHODS

### The GIGGLE Index

The GIGGLE index is based on a previously described temporal indexing method and consists of a set of B+Trees, one for each chromosome, represented among the database interval files. A B+Tree is a generalization of a binary tree where each node can have multiple keys, internal nodes contain only keys and facilitate tree searching, leaf nodes contain the key-value pairs, and adjacent leaf nodes are linked. Each key in an internal node is linked to two child nodes. The “left” link points to a node with keys that are less than the current key and the “right” link points to a node with keys that are greater than or equal to the current key. In the GIGGLE index, keys represent chromosomal positions, and the values associated with keys are lists of intervals that either start at that position (indicated by a “+” in Figure 1a and Supplemental Figure 1) or have ended just prior to that position (indicated by a “-” in Figure 1a and Supplemental Figure 1). Leaf nodes also contain a “leading” key (“L” in Supplemental Figure 1, omitted from Figure 1a) that stores intervals that start prior to the first key in the leaf, but have not ended by the last key in the prior leaf node (e.g., interval T1 in Supplemental Figure 1). While the leading values contain redundant information and require extra storage, they improve performance by preventing queries from having to load and scan other leaf nodes for intervals that start in some earlier leaf node (e.g interval T1).

### Bulk Indexing

To improve indexing efficiency, GIGGLE performs “bulk” indexing across many pre-sorted interval files (see **Data Format And Sorting Requirements**, Supplemental Figure 1a). In bulk indexing, a priority queue is used to is used to select the interval with the next lowest start position among the full set of files. The queue is loaded with one (the first) interval from each file and as intervals are removed from the queue the next interval from the corresponding file is added to the queue. For example, in Supplemental Figure 1b, Step 1 considers interval P1 from the Promoters file and Step 2 considers interval B1 from the TF binding sites file. Afre P1 is considered the next interval in the Promoters file (P2) is added to the queue. Similar in step 2 for B2 in the TF binding sites file.

Each interval is inserted into both the B+Tree based on its start position and an auxiliary priority queue that is keyed by the end coordinate (plus one). This priority queue is used to add intervals to the leading values and insert end positions into the B+Tree. If the start position of the current interval has been previously observed, then the interval start is added to the list of the existing value (intervals B2, P2, and T2 in Steps 4, 5, and 6 in Supplemental Figure 1b). Otherwise a new key is added to the current leaf for the start position (assuming the current leaf has not reached its maximum number of keys, which is set to 100 by default), and the interval start (“+”) is added to the new list that is associated with that key (intervals P1, B1, and T1 in Steps 1, 2, and 3 in Supplemental Figure 1b). Before a key is added for a new start position, all intervals with end-values less than or equal to the start value are removed from the priority queue and the interval ends (”-”) are either added to the lists of existing keys (interval B1 in Steps 4 in Supplemental Figure 1b) or new keys are created (intervals P1, T1, B2, P2, and T2 in Steps 4 and 6 in Supplemental Figure 1b). If at any point the current node becomes full, then a new leaf node is created, all intervals in the priority queue are added to the leading key value of the new node, and the new key is added to the new node (Step 4 in in Supplemental Figure 1b). Once all files have been processed and leaf node construction is complete, internal nodes are added by promoting the first key in each leaf node (other than the leftmost node) to a parent node (Step 7 in Supplemental Figure 1b). This process continues one level at a time until there is only one parent node.

### Searching the GIGGLE Index

A B+Tree search starts at the root node, proceeds down internal nodes, and terminates at a key in a leaf node. At each node in the search path, an internal search is performed among the keys in the node. While the current node is not a leaf node, the result of that search determines the next node in the path. If the matching key is less than or equal to the query, the path will follow the key’s the “right” link, otherwise it will follow the key’s “left” link. When the keys of a leaf node have been searched, the leaf and matching key are returned.

For a given query interval, GIGGLE performs a specialized range query to find overlapping intervals (the intersecting set) across all indexed files. First, the B+Tree is searched for the query interval’s start and end values, which gives the start leaf node and start key and the end leaf node and end key, respectively (Step 1 in Supplemental Figure 1c). Next, the intervals in the leading value of the start leaf node are added to the intersecting set (Step 2 in Supplemental Figure 1c). Then the keys in the leaf node are scanned from the first value up to and including the start key. At each key, starting intervals (“+”) are added to the intersecting set and ending intervals (“-”) are removed (Step 3 and 4 in Supplemental Figure 1c). Last, the remaining keys up to and including end key are scanned and the starting intervals are added at each key (Step 5 and 6 in Supplemental Figure 1c). If the start key does not equal the end key, then the search will use the links between leaf nodes.

### Data Format And Sorting Requirements

For indexing, GIGGLE supports VCF files (<u>https://samtools.github.io/hts-specs/VCFv4.3.pdf</u>) and BED files (<u>https://genome.ucsc.edu/FAQ/FAQformat.html#format1</u>) that have been sorted and bgzipped (<u>https://github.com/samtools/htslib</u>). Sorting is ascending lexicographical by chromosome, then ascending numerically by start and then by end. For searching, GIGGLE supports bgzipped VCF and BED files. These need not be sorted, but sorted files are likely to perform better due to cache performance. We provide a script in the GIGGLE repository (<u>https://github.com/ryanlayer/giggle/blob/master/scripts/sortbed</u>) that can sort and bgzip full directories using multiple processors. Otherwise the following command can be used on individual, uncompressed BED files:

~~~
LC_ALL=C sort −−buffer-size 2G -k1,1 -k2,2n -k3,3n **track.bed** | bgzip -c > track.bed.gz
~~~

## DATA SOURCES

### CHROMHMM / Roadmap Epigenomics data source

Tissue-based annotations were downloaded from:

<u>http://egg2.wustl.edu/roadmap/data/byFileType/chromhmmSegmentations/ChmmModels/coreMarks/jointModel/final/all.mnemonics.bedFiles.tgz</u>

These files were subsequently split and renamed into tissue/state-based files (e.g. Spleen/Enhancers). Detailed methods underlying this process can be found at: <u>https://github.com/ryanlayer/giggle/blob/master/examples/rme/README.md</u>

### UCSC Genome Browser data source

The full set of hg19 annotations were downloaded from: <u>http://hgdownload.cse.ucsc.edu/goldenPath/hg19/database</u>. The files with identifiable chromosome, start, and end values are converted to BED files. Detailed methods underlying this process can be found at: <u>https://github.com/ryanlayer/giggle/blob/master/examples/ucsc/README.md</u>

### MyoD ChIP-seq data source

ChIP-seq peaks from GSM1218850 are downloaded from: <u>ftp://ftp.ncbi.nlm.nih.gov/geo/samples/GSM1218nnn/GSM1218850/suppl/GSM1218850MB135DMMD.peak.tx</u> tgz

Peaks with a q-value greater than or equal to 100 are retained: Detailed methods underlying this process can be found at:

<u>https://github.com/ryanlayer/giggle/blob/master/examples/myod/README.md</u>

### GWAS variants for 39 autoimmune and non-autoimmune traits data source

A spreadsheet with a list of traits, chromosome, start, end, and other fields was downloaded from: <u>https://www.nature.com/nature/journal/v518/n7539/extref/nature13835-s1.xls</u> Detailed methods underlying this process can be found at:

### Fantom5 data source

The enhance expression matrix and associated metadata was downloaded from: <u>http://fantom.gsc.riken.jp/5/datafiles/latest/extra/Enhancers/Human.sample_name2libraryid.txt</u> <u>http://fantom.gsc.riken.jp/5/datafiles/latest/extra/Enhancers/human_permissive_enhancers_phase1and2e</u> <u>xpression count matrix.txt.gz</u>

Values were extracted from the matrix and placed in tissue-specific files. Detailed methods underlying this process can be found at:

<u>https://github.com/ryanlayer/giggle/blob/master/examples/fantom/README.md</u>

### Cistrome data data source

Reanalyzed ChIP-seq narrow peaks from raw GEO data was downloaded from: <u>http://cistrome.org/db/interface.htm</u>

With the following fields selected: Human_TF, Human_histone, Human_chromatin_accessibility, Human_other. Only peaks with a q-value greater than 100 were retained. For Figure 2d, all files had to pass two Cistrome CQ metrics: fraction of reads in peaks, and at last 500 peaks had to have 10 fold enrichment. We also removed files with less than 100 peaks with a q-value greater than or equal to 100. Detailed methods underlying this process can be found at:

<u>https://github.com/ryanlayer/giggle/blob/master/examples/cistrome/README.md</u>

## EXPERIMENTS

### Speed tests

Runtimes were for counting the number of intersection between a query set and a database set for GIGGLE (<u>https://github.com/ryanlayer/giggle</u>), BEDTOOLS (<u>https://github.com/arq5x/bedtools2</u>), and TABIX (<u>https://github.com/samtools/htslib</u>). The query sets had between 10 and 1 million 100 basepair intervals, and the databases were the ChromHMM predictions from Roadmap Epigenomics and the hg19 annotations from the UCSC genome browser. All tests were performed using a single core on the 2. 4GHz Intel Xeon processor (E5-2680 v4) with 25 MB of cache and a 510 MB/s read 485 MB/s write SSD drive (SM863a). Detailed methods underlying this process can be found at:

<u>https://github.com/ryanlayer/giggle/blob/master/experiments/speedtest/README.md</u>

### Relationship comparison

Two pairs of methods for quantifying the relationship between a query interval set and a database interval set were compared: the Fisher’s exact test of a 2x2 contingency table versus a Monte Carlo-base p-value and the odds ratio of a 2x2 contingency table versus a Monte Carlo-base enrichment. The query set was the GWAS variants associated with Crohn’s disease, and the database was the ChromHMM predictions from Roadmap Epigenomics. Monte Carlo simulations were performed using BITS (<u>https://github.com/arq5x/bits</u>), a simulation was performed for the intersection of the GWAS variants and each tissue/genomic state interval set, and each simulation consisted of 1,000 rounds. Detailed methods underlying this process can be found at:

<u>https://github.com/ryanlayer/giggle/blob/master/experiments/mcvstable/README.md</u>

### MyoD heat map

The GIGGLE scores for MyoD ChIP-seq peaks searched against the ChromHMM predictions from Roadmap Epigenomics. Only the peaks with a q-value greater that 100 were used. The cell line and tissue names are in Supplementary File 1. Detailed methods underlying this process can be found at: <u>https://github.com/ryanlayer/giggle/blob/master/experiments/chipseq/README.md</u>

### Crohn’s disease heat map and Autoimmune / non-autoimmune heat map

The GIGGLE scores for the sets of GWAS variants associated with Crohn’s disease and other traits were searched against the ChromHMM predictions from Roadmap Epigenomics. For Figure 2c, the left two columns correspond to the scores from the Enhancer state from ChromHMM for each tissue and the right two columns correspond to the scores from the Strong Transcription state. Within these major columns the left minor column corresponds to the autoimmune disorders and the right minor column to the non-autoimmune traits. The categorization of these traits was retained from Farh *et al^11^.* The cell line and tissue names are in Supplementary File 1. The autoimmune disorders and the non-autoimmune traits are listed in Supplemental File 1. Detailed methods underlying this process can be found at:

<u>https://github.com/ryanlayer/giggle/blob/master/experiments/gwas/README.md</u>

### Cistrome ER

The GIGGLE scores for the ChIP-seq peak files of ERS1 from the MCF-7 cells line that passed quality control (described above) were searched against all other MCF-7 cell line results that also passed quality control. Only the peaks with a q-value greater that 100 were used. The full set of accession numbers is Supplementary File 1. Detailed methods underlying this process can be found at: <u>https://github.com/ryanlayer/giggle/blob/master/experiments/cistrome/README.md</u>

### Cistrome MCF-7

The GIGGLE scores for all ChIP-seq peak files from the MCF-7 cell line that passed quality control were searched against themselves. Only the peaks with a q-value greater that 100 were used. The full set of accession numbers is Supplementary File 1. Detailed methods underlying this process can be found at: <u>https://github.com/ryanlayer/giggle/blob/master/experiments/cistrome/README.md</u>

## GIGGLE COMMAND LINE AND PROGRAMMING INTERFACES

### Command line interface

#### Indexing

~~~
giggle index -i “intervals/*.bed.gz” -o interval_index -s
~~~

#### Searching.

~~~
giggle search -i interval_index -r chr1:1000000-2000000
giggle search -i interval_index -q query.bed.gz
~~~

### C interface

#### Indexing.

~~~
#include                                 “giggle_index.h”
int main(int argc, char **argv) {
 uint64_t num_intervals = giggle_bulk_insert(“intervals/*.bed.gz”, “interval_index”, 1);
 return 0;
}
~~~

#### Searching.

~~~
#include                                 “giggle_index.h”
int main(int argc, char **argv) {
 struct giggle_index *gi = giggle_load(“interval_index”, block_store_giggle_set_data_handler);
struct giggle_query_result *gqr = giggle_query(gi, “chr1”,1000000, 2000000, NULL);
uint32_t i;
 for(i = 0; i < gqr–>num_files; i++) {
  struct file_data *fd = file_index_get(gi–>file_idx, i);
  if (giggle_get_query_len(gqr, i) > 0)) {
   char *result;
   struct giggle_query_iter *gqi = giggle_get_query_itr(gqr, i);
   while (giggle_query_next(gqi, &result) == 0)
     printf(“%s\t%s\n”, result, fd−>file_name);
  giggle_iter_destroy(&gqi);
  }
 }
 giggle_query_result_destroy(&gqr);
 giggle_index_destroy(&gi);
 return 0;
}
~~~

**Python interface**: <u>https://github.com/brentp/python-giggle</u>

#### Indexing.

~~~
from giggle import Giggle
index = Giggle.create(‘interval_index’, ‘intervals/*.bed.gz’)
~~~

#### Searching.

~~~
from giggle import Giggle
index = Giggle(‘interval_index’)
print(index.files)
result = index.query(‘chr1’, 9999, 20000)
print(result.n_files)
print(result.n_total_hits)
print(result.n_hits(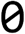))
for hit in result[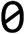]:
 print(hit) # hit is a string
~~~

**Go interface**: <u>https://github.com/brentp/go-giggle</u>

#### Indexing.

~~~
import (
 giggle “github.com/brentp/go-giggle”
 “fmt”
)
func main() {
 index := giggle.New(“interval_indexr”, “intervals/*.bed.gz”)
}
~~~

#### Searching.

~~~
import (
 giggle “github.com/brentp/go-giggle”
 “fmt”
)
func main() {
 index := giggle.Open(“interval_index”)
 res := index.Query(“1”, 565657, 567999)
 // all files in the index
 index.Files()
 // int showing total count
 res.TotalHits()
 // []uint32 giving number of hits for each file
 res.Hits()
 var lines []string
 # access results by index of file.
 lines = res.Of(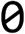)
 fmt.Println(strings.Join(lines, “\n”))
 lines = res.Of(1)
}
~~~

## SUPPLEMENTARY TABLES

**Supplementary Table 1.** Top GIGGLE results for MyoD ChIP-seq peaks against the CHROMHMM Roadmap data set.

| Roadmap Cell Type / Genomic State | GIGGLE Score | Odds Ratio | Fisher's exact test |
| --- | --- | --- | --- |
| HSMM Skeletal Muscle Myoblasts / Enhancers | 1205.91 | 65.34 | 1.02e-200 |
| HSMM Skeletal Muscle Myotubes / Enhancers | 1188.27 | 60.23 | 1.05e-201 |
| HSMM Skeletal Muscle Myotubes / Flanking Active TSS | 1188.08 | 60.35 | 1.415e-201 |
| HSMM Skeletal Muscle Myoblasts / Flanking Active TSS | 1136.93 | 49.98 | 3.39e-202 |
| Fetal Muscle Leg / Enhancers | 1008.06 | 32.45 | 1.56e-201 |

**Supplementary Table 2.** Top GIGGLE results for MyoD ChIP-seq peaks against the FANTOM5 data set.

| FANTOM5 Cell Type | GIGGLE Score | Odds Ratio | Fisher's exact test |
| --- | --- | --- | --- |
| Skeletal muscle cells differentiated into Myotubes donor 1 | 311.51 | 40.97 | 7.03e-59 |
| Skeletal Muscle Cells donor 1 | 188.26 | 28.90 | 1.614e-39 |
| Myoblast differentiation to myotubes day 10 donor 1 | 154.52 | 32.22 | 1.44e-31 |
| Myoblast donor 3 | 146.10 | 29.67 | 1.34e-30 |
| Myoblast differentiation to myotubes day 2 donor 3 | 137.15 | 10.51 | 3.80e-41 |

**Supplementary Table 3.** Top GIGGLE results for Crohn’s disease GWAS SNPs against the CHROMHMM Roadmap data set.

| Roadmap Cell Type / Genomic State | GIGGLE Score | Odds Ratio | Fisher's exact test |
| --- | --- | --- | --- |
| Primary T CD8+ naive cells / Enhancers | 154.68 | 6.68 | 3.55e-57 |
| Primary T helper cells / Enhancers | 115.32 | 5.63 | 5.616e-47 |
| Primary T CD8+ memory cells / Enhancers | 113.43 | 6.19 | 7.45e-44 |
| Fetal Thymus / Enhancer | 113.09 | 5.92 | 8.54e-45 |
| Primary T helper cells / Enhancers | 112.8 | 5.81 | 3.82e-45 |

**Supplementary Table 4.**
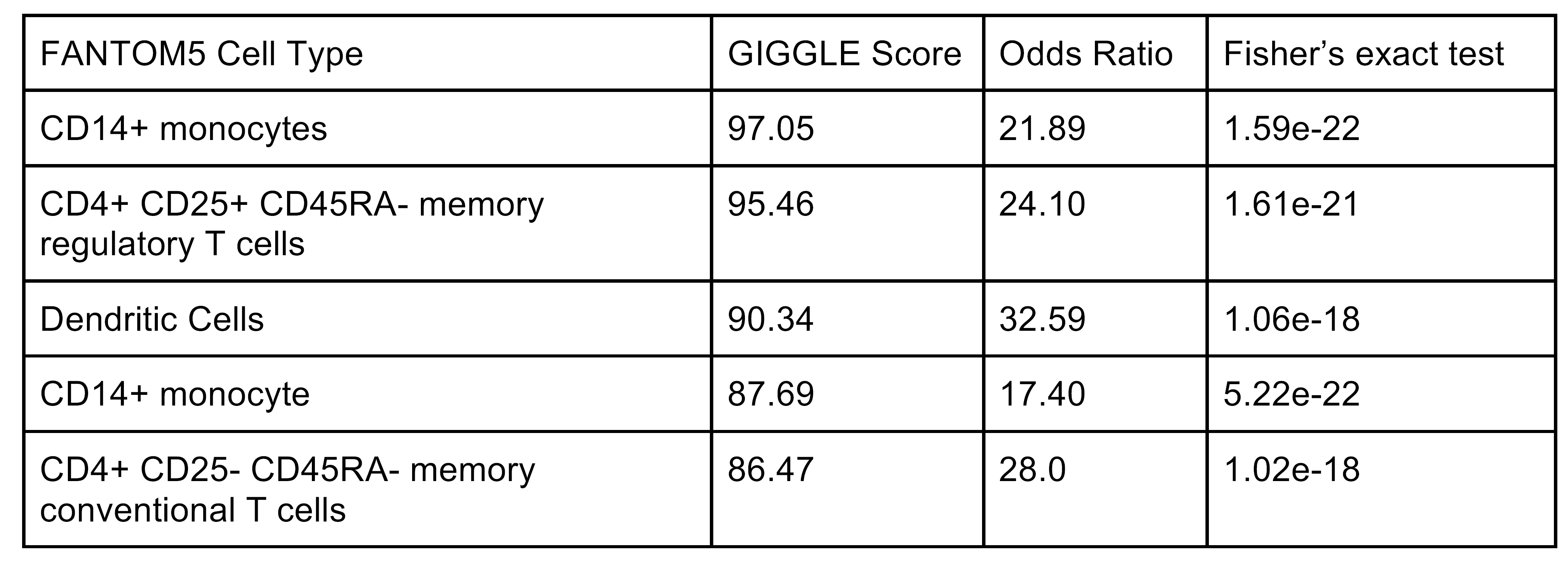
Top GIGGLE results for Crohn’s disease GWAS SNPs against the FANTOM5 data set.

## SUPPLEMENTARY FIGURES

**Supplemental Figure 1.**
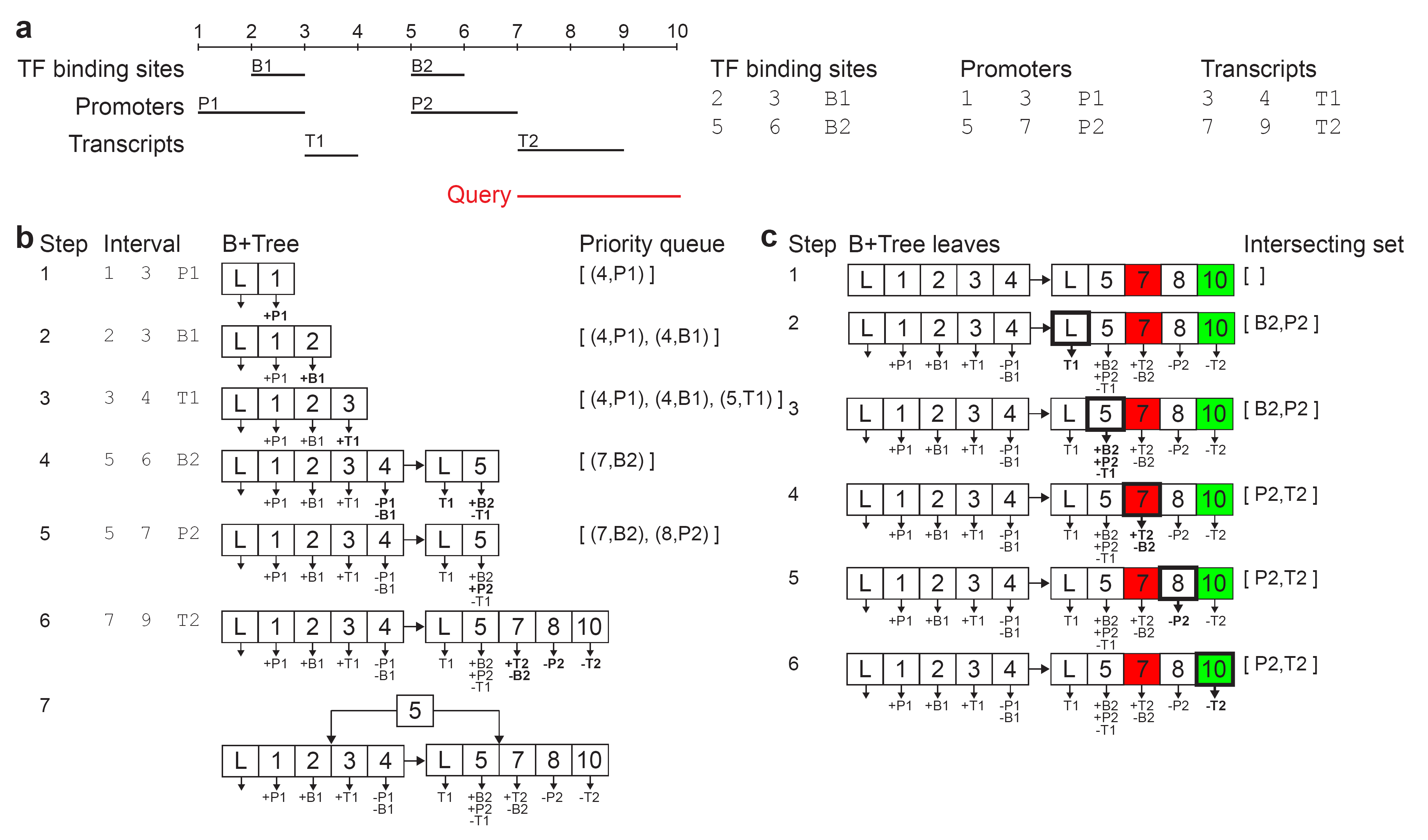
(**a**) Three example annotation sets shown graphically (left) and encoded in files (right) by start position, end position, and ID. (**b**) GIGGLE’s bulk indexing process. (**c**) The GIGGLE interval search process.

**Supplemental Figure 2.**
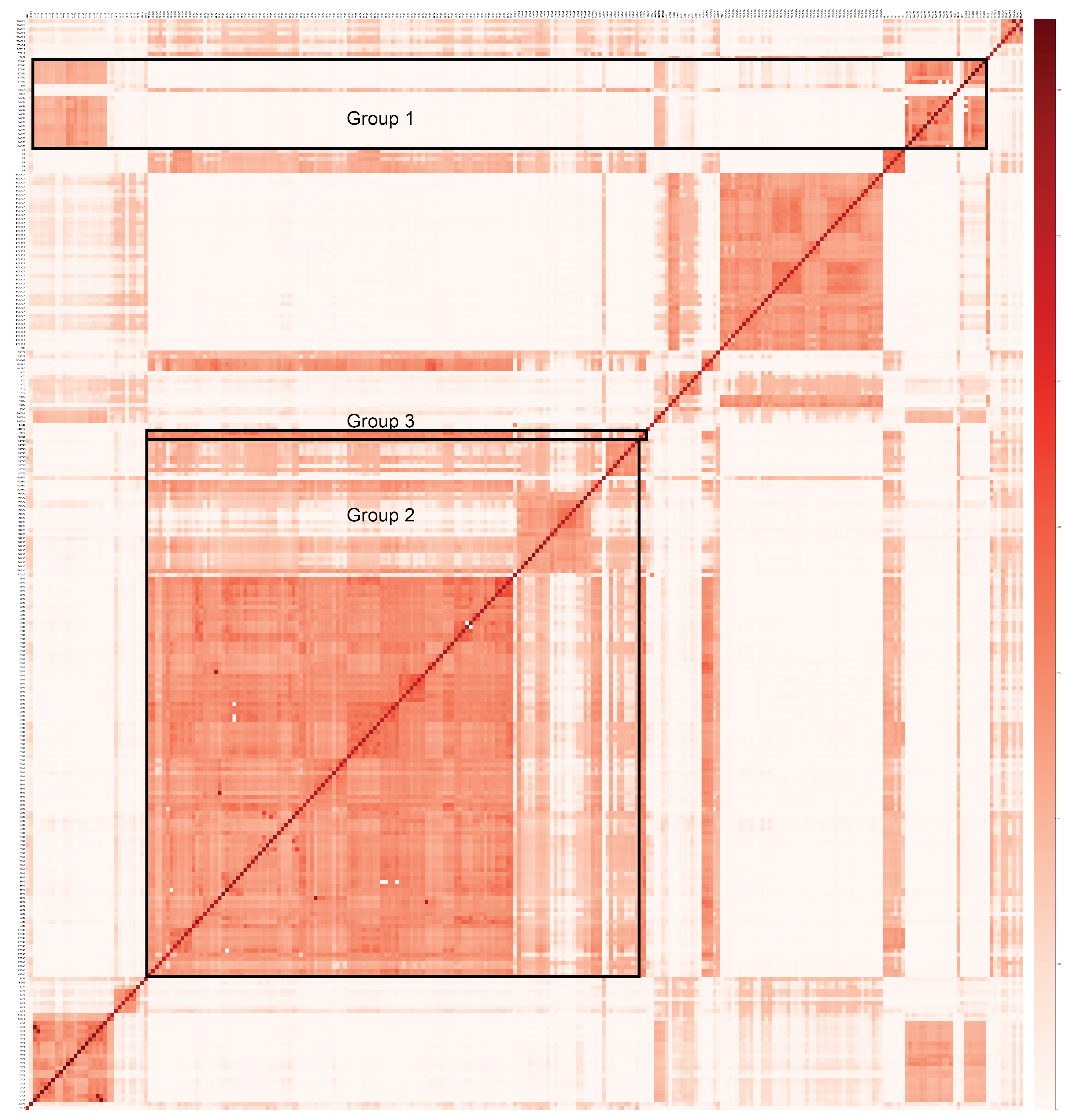
The GIGGLE scores for all pairwise combinations of the ChIP-seq datasets for the MCF-7 cell line. Group 1 highlights the relationship between CTCF, RAD21, and STAG1. Group 2 highlights ERS1, FOXA1, GATA3, and EPS300. Group 3 shows an unexpected relationship between H2AFX and GREB1.

**Supplemental Figure 3.**
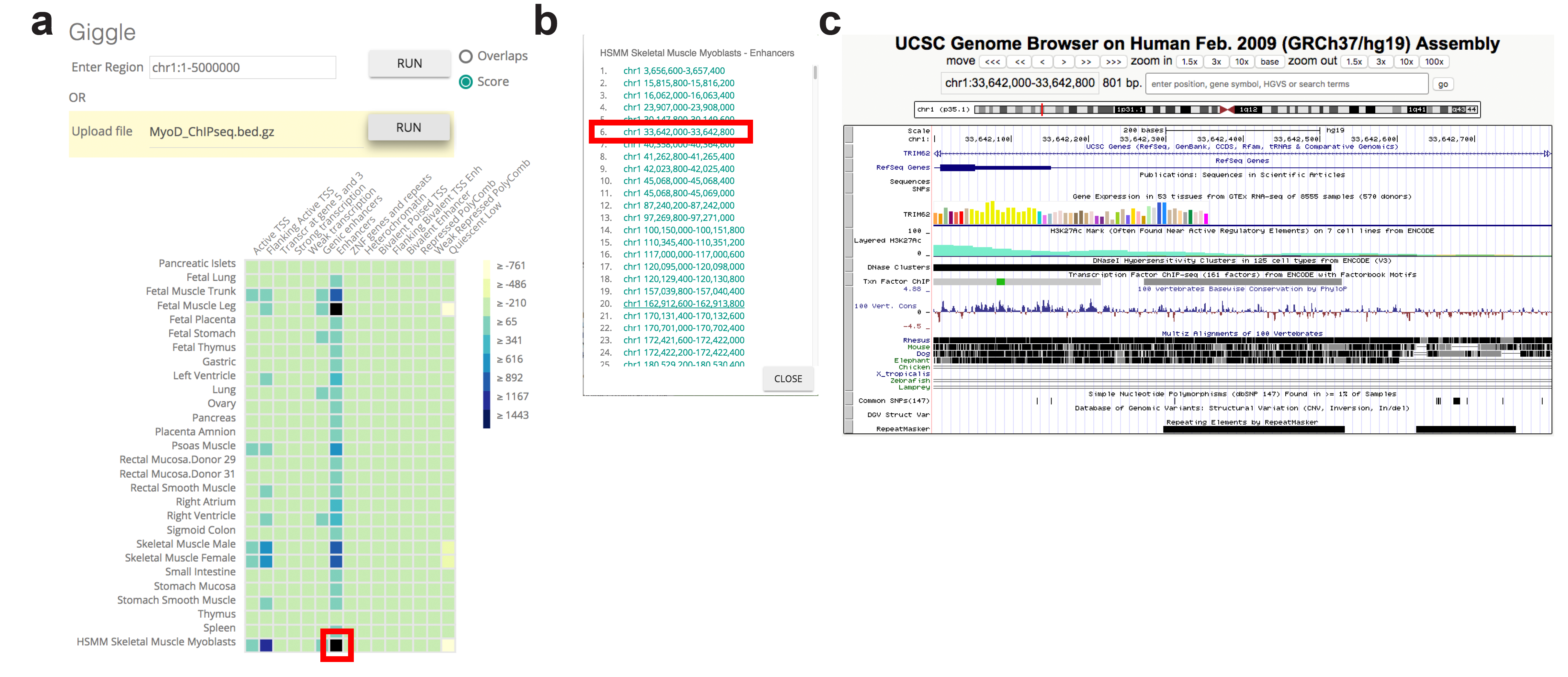
A web interface that integrates data from of Roadmap and the UCSC genome browser. (**a**) Users specify either a single interval or file to upload as the query, and the server responds with the GIGGLE results from an index in a heatmap. In this case the index is of CHROMHMM prediction from Roadmap. The color of each cell indicates the GIGGLE score, and users can click on a cell (e.g., Myoblast enhancers, marked in red) for more information. (**b**) When a cell is selected by the user, a window opens that contains the list of intervals in that particular Roadmap cell type/genome state annotation that overlap the query. Each interval is a link that can be followed (e.g., chr1:33642000-33642800, marked in red) for more information. (**c**) When an interval is selected, that interval becomes a query to a GIGGLE index of the UCSC genome browser tracks. The result gives the set of tracks that contain an interval that overlaps the query, and the web interface opens a window with a “smartview” where only those tracks with overlaps are displayed.

